# unitas: the universal tool for annotation of small RNAs

**DOI:** 10.1101/165175

**Authors:** Daniel Gebert, Charlotte Hewel, David Rosenkranz

## Abstract

**Background:** Next generation sequencing is a key technique in small RNA biology research that has led to the discovery of functionally different classes of small non-coding RNAs in the past years. However, reliable annotation of the extensive amounts of small non-coding RNA data produced by high-throughput sequencing is time-consuming and requires robust bioinformatics expertise. Moreover, existing tools have a number of shortcomings including a lack of sensitivity under certain conditions, limited number of supported species or detectable sub-classes of small RNAs.

**Results:** Here we introduce unitas, an out-of-the-box ready software for complete annotation of small RNA sequence datasets, supporting the wide range of species for which non-coding RNA reference sequences are available in the Ensembl databases (currently more than 800). unitas combines high quality annotation and numerous analysis features in a user-friendly manner. A complete annotation can be started with one simple shell command, making unitas particularly useful for researchers not having access to a bioinformatics facility. Noteworthy, the algorithms implemented in unitas are on par or even outperform comparable existing tools for small RNA annotation that base on available ncRNA sequence information.

**Conclusions:** unitas brings together annotation and analysis features that hitherto required the installation of numerous different bioinformatics tools which can pose a challenge for the non-expert user. With this, unitas overcomes the problem of read normalization. Moreover, the high quality of sequence annotation and analysis, paired with the ease of use, make unitas a valuable tool for researchers in all fields connected to small RNA biology.

## Background

Small non-coding (snc-) RNAs are important players in diverse cellular processes, often acting as guide molecules in transcriptional and post-transcriptional gene regulation [l, 2, 3]. Micro (mi-) RNAs, short interfering (si-) RNAs and Piwi- interacting (pi-) RNAs constitute their most prominent representatives but the number of described sncRNA classes continuously increases. Moreover, degradation products of larger RNA molecules such as rRNA or tRNA fragments further contribute to sequence heterogeneity of sncRNA transcriptomes [4, 5]. As diverse as their source molecules are the places where sncRNAs can be found within an organism, ranging from nuclear and cytoplasmic localization inside a cell, to extra-cellular exosomes being released into diverse body fluids [6, 7]. Studying the role of sncRNAs in diverse biological contexts typically involves high-throughput sequencing of sncRNAs derived from total RNA extracts. Subsequent disentangling of the complex composition of such sncRNA transcriptomes is one of the initial steps in sequence data processing and critical for all kinds of downstream analysis. As the use of high throughput sequencing technologies becomes more and more common, while this does not necessarily apply to bioinformatics knowhow, a robust and easy to use solution for reliable annotation of sncRNA sequence datasets is highly desirable.

So far, annotation of sncRNA sequence datasets is demanding for various reasons. On the technical side, existing tools cover particular aspects of sequence annotation (e.g. miRNA annotation) which means that complete annotation including all types of sncRNAs requires installation of a set of programs with different dependencies, some of which are restricted to specific operating systems. lllustrating the complexity of the task, a typical annotation process could include the following steps: i) 3’ adapter recognition with Minion [8] or DNApi [9], ii) adapter trimming with e.g. reaper [8] or cutadapt [10], iii) filtering of low complexity sequences with dustmasker [11] or RepeatSoaker [12], iv) miRNA annotation with Chimira [13], v) annotation of tRNA-derived fragments with tDRmapper [14] or MlNTmap [15], vi) annotation of other ncRNA or mRNA fragments with NCBl BLAST and, if applicable, vii) annotation of phased RNAs with PhaseTank [16] or viii) annotation of putative piRNAs by mapping sncRNA sequences to known piRNA producing loci [17]. However, when having established a local annotation pipeline it is almost impossible to correctly normalize the obtained results in case that a given sequence maps to different types of non-coding RNA. Even with a profound bioinformatics expertise, custom annotation is challenging due to the fact, that reference non-coding RNA sequences are stored at different online databases such as Ensembl database, miRBase, GtRNAdb and SlLVA rRNA database. Further, mapping sncRNA sequences to reference sequences, once having gathered a complete collection, and subsequent parsing of the obtained results is bedeviled by, e.g., the presence of isomiRs or post-transcriptionally adenylated or uridylated miRNAs.

In order to facilitate and speed-up sncRNA annotation while making the obtained results comparable across different studies, we have developed unitas, a tool for sncRNA sequence annotation that requires not more than a computer with internet connection. Our aim is to provide a maximally convenient tool that runs with an absolute minimum of prerequisites on any popular operating system, making high-quality sequence annotation available for everyone. By providing complete annotation with one tool we intend to tackle the problem of normalization of multiple mapping sequences. ln addition, we designed all annotation and analysis algorithms with the aim to overcome a number of limitations of existing tools, in order to make unitas the means of choice compared to a notional pipeline with state-of-the-art tools connected in series. The unitas source code and precompiled executable files are freely available at http://www.smallrnagroup. uni-mainz.de/software.html.

## lmplementation

### General requirements

We provide precompiled standalone executable files of unitas for Linux, Mac and Windows systems. unitas itself is written in Perl and designed to run with an absolute minimum of prerequisites, relying on Perl core modules, or modules which are part of widely used free Perl distributions such as Archive:: Extract and LWP:: Simple. Perl is commonly preinstalled on Linux and MacOS systems, where users can run the unitas Perl script without any further requirements. Windows users that prefer to run the Perl script rather than the executable file may have to install a free Perl distribution such as ActivePerl or Strawberry Perl. More detailed information and help is available in the unitas documentation. Since unitas uses publicly available online databases for sncRNA annotation, the program needs an internet connection when run for the first time. Later runs can use previously downloaded data. lnput files can be sequence files in FASTA or FASTQ format (with or without 3’ adapter sequence), or alternatively map files in SAM or ELAND3 format. Some data analysis features are only available when using map files as input.

### Reference sequence data management

Sequence annotation with unitas relies on publicly available reference sequences from Ensembl [18], miRBase [19], GtRNAdb [20], SlLVA rRNA database [211] and piRNA cluster database [17] (Fig. l). Currently, unitas supports 835 different species or strains for which information on ncRNAs is available at least in one of the Ensembl databases. Prior to annotation, unitas downloads a collection of latest reference sequences which are stored in a separate folder on the local machine for subsequent mapping. As availability of reference sequences is crucial, unitas is designed to address possible challenges that can occur during acquisition of that data. Since database URLs often change with new releases or updates of reference sequences, relying on URLs stored inside the programs source code would require frequent updates of the unitas software itself. Therefore, unitas connects to the Mainz University Server (MUS) and loads the latest list of URLs for downloading the required reference sequence data. However, in the event of these URL not being up to date (URLs are updated monthly), unitas ultimately downloads the required sequence data directly from MUS where the datasets are available via stable URLs and are synchronized regularly (Fig. l). By default, downloaded sequence datasets are used for subsequent unitas runs without anew downloading by default. Users can also download reference sequence data collections for any supported species at any time and use the downloaded data for later offline runs. The downloaded sequence data can be updated anytime.

**Fig. 1:**
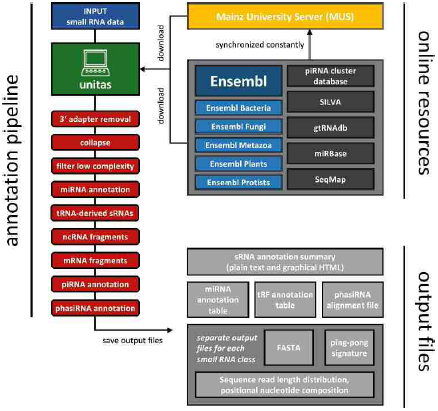
Architecture of the unitas workflow ata glance.

### Automated 3’ adapter recognition and trimming

Standard cloning protocols for small RNA library preparation prior to high throughput sequencing involve ligation of adapter molecules to both ends of a RNA molecule. During sequencing, the sequencing primer typically hybridizes to the 3’ end of the 5’ adapter, which means that the resulting sequence read starts with the original small RNA sequence and, given a sufficient read length, ends with the 3’ adapter sequence. However, there exist manifold commercially available 3’ adapters that can be used for library construction. ln addition, these adapters can contain different index/barcode sequences. Finally, working groups may even use custom made adapter molecules. Since information on 3’ adapter sequences that were used to generate a small RNA dataset is not always available or at least difficult to find out, we integrated an adapter recognition and trimming module that can be applied using the option ‘-trim’.

Initially, unitas identifies the most frequently occurring sequence ignoring sequence read positions 1 to 22 (typical length for miRNAs). unitas Adjusts The Length of the motif, *m*, to be identified automatically according to the formula *m* = *n*-22 (as long as:6 ≤ m ≤ 12) where *n* refers to the sequence read length. A first round of adapter trimming is the performed based on the identified motif allowing 2 mismatches for 12 nt motifs, 1 mismatch formotifs ≤ 11nt and 0 mismatch for motifs ≤ 8 nt. If the original motif is not found with inagivens equenceread, unitas truncates the motifs sequentially by one 3’nt and checks for it occurrence at the very 3’. end of the sequence read until the motifis found or the motif length falls below 6 nt. Following this first round of adapter trimming, unitas checks the positional nucleotide composition of the trimmed sequence reads and will remove further 3’. nucleotide positions in case they exceed a specified nucleotide bias (default = 0.8). Noteworthy, there may exists cenariosin which unitas will not detect the correct 3’. adapters equences when using the default settings, particularly in cases with short library read length (≤35 nt) combinedwith a highamount of reads that share 3’. similarity such as, e.g., tRNA derived fragments. In these special cases, adapter recognition can be improved by increasing the amount of 5’.positions to beignored when searching for frequent sequence motifs(option :trim-ignore-5p[n]).

### Filtering low complexity reads

To filter out low complexity reads, unitas employs an advanced version of the duster algorithm from the NGS TOOLBOX [22]. By default, sequence reads with a length faction *f* > 0.75 being composed of one repetitive sequence motif (default motif length= l-5 nt) are rejected. Further, sequences with a length fraction

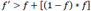

being composed of only two specific nucleotides are also rejected.

### miRNA annotation

Unitas performs miRNA annotation in several consecutive steps. Mature miRNA sequences and miRNA hairpin sequences are downloaded from miRBase and miRNAs are annotated in the following order: i) Canonical miRNAs of the species in question, ii) post-transcriptionally 3’-tailed canonical miRNAs of the species in question, iii) offset miRNAs of the species in question, iv) post-transcriptionally 3’-tailed offset miRNAs of the species in question. Subsequently, this procedure is repeated using miRNA sequence data from all other species included in miRBAse, which is particularly useful for those species with bad miRNA annotation status considering the fact that many miRNA sequences are widely conserved. Since the according output file comprises information on the source species of each matched miRNA gene, unitas users are able to assess the relevance of each match in a case-dependent manner. However, it is important to be aware that this approach will not identify lineage-specific miRNA genes, which can only be identified using *de novo* prediction tools. Nevertheless, accurate filtering of known miRNA sequences will be helpful for downstream *de novo* miRNA prediction. By default, the maximum number of allowed non-template 3’ nucleotides is 2 and the maximum number of allowed internal modifications is l. ln order to map sncRNA sequences to miRNA precursors (or to other ncRNA sequences in later annotation steps), unitas employs the mapping tool SeqMap [23] which not requires prior indexing of reference sequences and allows subsequent analysis of non-template 3’-nucleotides.

### ncRNA/mRNA annotation

Following miRNA annotation, sequences that do not correspond to miRNAs are mapped to a species-specific collection non-coding RNA and cDNA sequences downloaded from Ensembl database [18], Genomic tRNA database [20] and SlLVA rRNA database [21]. Read counts of sequences that match different classes of reference sequences equally well are apportioned according to the simple equation:

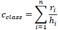

where cclass refers to the read counts for ncRNA/cDNA class, while *n* is the total number of non-identical input sequences that map to this class and *ri* and *hi* refer to read counts and hits to different ncRNA/cDNA classes of input sequence *i*, respectively. During this process, special attention is payed to sequence reads matching tRNAs since different classes of functional tRNA derived fragments, so-called tRFs, have been described in the recent past [24, 25, 26, 27, 28, 29, 30]. unitas classifies these sequences into 5’ tRFs (5’ to D-loop), 5’ tR-halves (5’ to Anticodon-loop), 3’ tRFs (Tψ C-loop to 3’), 3’ CCA-tRFs (Tψ C-loop to 3’CCA), 3’ tR-halves (Anticodon-loop to 3’), tRF-l (3’ end of mature tRNA to oligo-T signal), tRNA-leader (sequence upstream of 5’ ends of mature tRNAs) and misc-tRFs (miscellaneous tRFs). Worth mentioning, unitas relies on available ncRNA annotation and will not perform *de novo* prediction of ncRNA genes that encode, e.g., tRNAs or rRNAs.

### piRNA annotation

Considering the fact that piRNAs are highly diverse and virtually not conserved across different species, piRNA annotation based on sequence is challenging. However, many piRNAs originate from few genomic loci, many of which are annotated in the piRNA cluster database [17]. Providing that information on piRNA clusters is available for the species in question, sequences that were not annotated as (fragment of) any other class of non-coding RNA are mapped to known piRNA producing loci of the respective species. Since almost every nucleotide position within a piRNA precursor transcript can give rise to the 5’ end of a mature piRNA, though there is certainly a bias for 5’-U, this procedure more reliably identifies putative piRNAs compared to the approach of directly mapping sequence reads to annotated piRNAs. Further evidence for the presence of genuine piRNAs can be obtained from sequence read length distribution and positional nucleotide composition which unitas outputs for each class of small RNAs separately. Providing that the input file provided by the user represents a map file, unitas can further screen the map file for the so-called ping-pong signature (using the option -pp), which refers to a bias for l0 nt 5’ overlaps of mapped sequence reads which arises from secondary piRNA biogenesis (ping-pong cycle) and indicates the presence of primary and secondary piRNAs. Screening for a ping-pong signature also includes calculation of a Z-score according to the method described by Zhang and coworkers [31].

### phasiRNA annotation

The commonly applied m ethod for identification of phased RNAs bases on calculation of a so-called phase score, *P*. After consolidation of mapped reads from both strands with an offset of 2 nt for minus strand mapped reads, *P* results from the following formula:

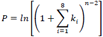

in which *n* refers to the number of phase cycle positions occupied by at least one small RNA read within an eight cycle window, and *K* refers to the total number of reads for all small RNAs with Consolidated start coordinates in a given phase within an eight cycle window [32].

Although the given formula yields higher *P* values with Increasing *k* or *n*, the weighting between both factors, and finally the decision of which threshold to choose For *P* is rather arbitrary. We therefore decided to use a different method, which utilizes the binomial distribution To calculate the probability *p* to observe a defined number (or more) of phased reads within a Given sliding window (defaul = 1kb) according to the formula:

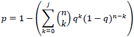

in which *j* refers to the observed number of reads with Length *I* in a specified phase *n* refers to the total number of reads with length *i* and *q* is given by l/*i* and refers to the probability of a read to be located in a given phase, assuming that a sequence read can map to any position within the sliding window with equal probability. As is the case for calculation of *P*, reads mapped to different strands are consolidated prior to calculation of *p*. lf the *p* value of a locus under examination is below the critical value (default = 0.05, with strict Bonferroni correction based on the number of analyzed sliding windows), unitas applies further thresholds to reduce the rate of false positive predictions. By default, the fraction of phased RNAs has to be ≥ 50% of all mapped reads within a sliding window. Further, the phased reads must map to ≥ 5 different loci while not more than 90% of the phased reads must derive from one strand. Critical values for each of the mentioned parameters, including *p* and sliding window size can be adjusted by the user. Prediction of phasiRNAs requires map files (SAM or ELAND3) as input and can be performed with the option ‘-phasi [*n*]’ where n refers to the length of the phased RNAs.

## Results

We have tested unitas using a number of artificial datasets, real RNA-seq data and combinations of both. A detailed description of the datasets and the methods that were applied to generate them can be found in Additional file l (Supplementary Methods).

### 3’ adapter identification and trimming

The first steps in the analysis of small RNA data usually involve the removal of sequencing adapters from 3’ ends, for which numerous tools exist. However, this task becomes problematic if the adapter sequence is not known, e.g. if a dataset is deposited without the appropriate information. The number of programs for adapter prediction, in contrast to removal, is rather limited. The only published tools for this purpose are DNApi [9] and Minion from the Kraken package [8], which also contains Reaper for adapter trimming.

To test the efficiency of the 3’ adapter identification and trimming function of unitas, we processed ten randomly chosen datasets from the NCBl Sequence Read Archive and put the perfomance into comparison to the existing software. Both, unitas and DNApi, reliably predicted the correct adapter sequences in all cases, whereas Minion predicted a false adapter with a slightly deviated sequence for one of the ten libraries (SRA accession: SRR5l30l42), leading to a considerably reduced efficacy in subsequent read trimming by Reaper (Fig. 2a). Altogether, in eight instances unitas removed more adapter sequences than Reaper, hence resulting in higher quantities of trimmed reads that could be mapped perfectly to the corresponding genome (+ 9.7% on average, Additional file 2: Table Sl).

**Fig. 2:**
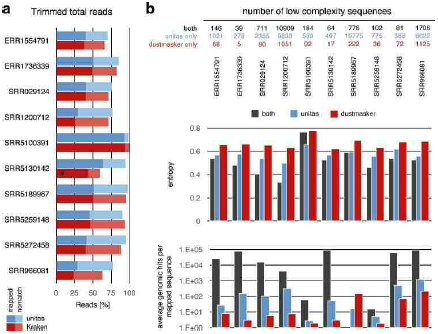
Adapter trimming and removal of low complexity reads. a Efficiency of 3’ adapter identification and trimming by unitas and Kraken tool kit. Asterisk marks the dataset for which adapter prediction by Minion failed. Trimmed reads were mapped to the corresponding genome without mismatches. b Removal and analysis of low complexity sequences filtered by unitas and dustmasker.

### Removal of low complexity sequences

The presence of low complexity reads can weaken biological signals within RNA-seq datasets and it has been demonstrated, that the correlation between RNA-seq and microarray gene expression data can be improved with strict filtering of sequences that map to genomic regions with low sequence complexity [12]. Since this method relies on the availability of a RepeatMasker annotation for the genome in question, we implemented a low complexity filter upstream of sequence annotation. We compared our filter with dustmasker, a popular tool for masking low complexity regions in DNA sequences, which is part of the NCBl blast+ package [1l]. We ran dustmasker on ten adapter-trimmed NGS sequence datasets (see above) with default settings and discarded sequence reads with more than 75% of bases being masked to produce results that are comparable to the results generated by unitas, which by default filters sequences being composed of more than 75% of repetitive sequence motifs.

First, we found that unitas generally filters more sequences (Fig. 2b top), which taken by itself is certainly not indicative to favor one tool over the other since both algorithms can easily be adjusted to filter more or less sequences by changing the corresponding thresholds. Therefore, we in-depth analyzed the complexity of sequences filtered by both tools as well as those sequences that were filtered either by unitas or by dustmasker. We quantified the complexity of filtered sequences based on sequence entropy [33], Wootton-Federhen-complexity [34] and gzip (developed by Jean-loup Gailly and Mark Adler) compression ratio using the program SeqComplex, which was written by Juan Caballero. As expected, sequences filtered by both tools usually exhibit the lowest degree of entropy. Further, sequences filtered only by unitas exhibit a lower degree of entropy compared to sequences filtered only by dustmasker, in spite of the fact that unitas filters more sequences (Fig. 2b middle). According to Wootton-Federhen-complexity and gzip compression ratio, sequences filtered only by unitas also exhibit lower complexity compared to sequences filtered only by dustmasker and even compared to those sequences filtered by both tools (Additional file 3: Table S2). We further wanted to check whether these rather theoretical assessments can be translated into a biological dimension. To this end we mapped the filtered sequences to the respective genomes and counted the number of genomic hits per sequence, assuming that the amount of information obtained by mapping a specific sequence decreases with a growing number of genomic hits. ln line with the previous results, sequences filtered by both tools show the highest number of genomic hits, thus providing the lowest amount of information (Fig. 2b bottom). With one exception, sequences filtered exclusively by unitas map more frequently to the genome compared to sequences filtered exclusively by dustmasker (Fig. 2b bottom). Together, these results demonstrate that unitas filters sequences with low complexity in a more sensitive and more specific manner.

### Annotation of miRNAs

Numerous programs for miRNA annotation in small RNA-seq data have been published in the past with varying focuses [35, 36, 37]. To compare the performance of unitas on this task we chose Chimira, which is a recent tool with a similar range of functions, primarily aiming at miRNA expression and modification analysis [13]. Chimira is a web-based system, accepting multiple input files in FASTA or FASTQ format at once and supporting 209 genomes so far. lnput reads are mapped against miRBase [19] hairpin sequences using BLASTn with two tolerated mismatches, identifying modifications at 3’ and 5’ ends, aswell as internal substitutions(single nucleoti polymorphisms, SNPs) and ADAR-dependent editing events in the process. However, in order for an internal modification to be classified as a SNP, an arbitrary value of 70% is applied as a threshold for the ratio of modification counts to overall counts. For a controlled comparison, we produced an artificial miRNA dataset based on human hairpin sequences from miRBase (release 2l), incorporating internal modifications and 3’ tailings. Of overall 466,8l0 generated reads, unitas identified 99.9% as miRNAs, while Chimira detected only 85.8% of the original set. Moreover, unitas showed higher precision in assigning read counts to respective miRNA genes of origin than Chimira did, indicated by Pearson correlation coefficients of 0.95l4 and 0.9l46, respectively (Fig. 3a, Additional file 4: Table S3). Furthermore, unitas detected 3’ tailings and internal modifications more reliably, whereas Chimira barely showed the latter type, probably due to the considerably high (70%) threshold for internal modifications (Fig. 3b). Noteworthy, the test dataset was designed to include all possible combinations of offset-, tailing-, and mismatch-scenarios without any weighting between canonical and non-canonical sequences. Consequently, the differences between unitas and Chimira annotations are typically less marked for real biological datasets (these observations are principally true for annotation of tRNA fragments as well, see below).

**Fig. 3:**
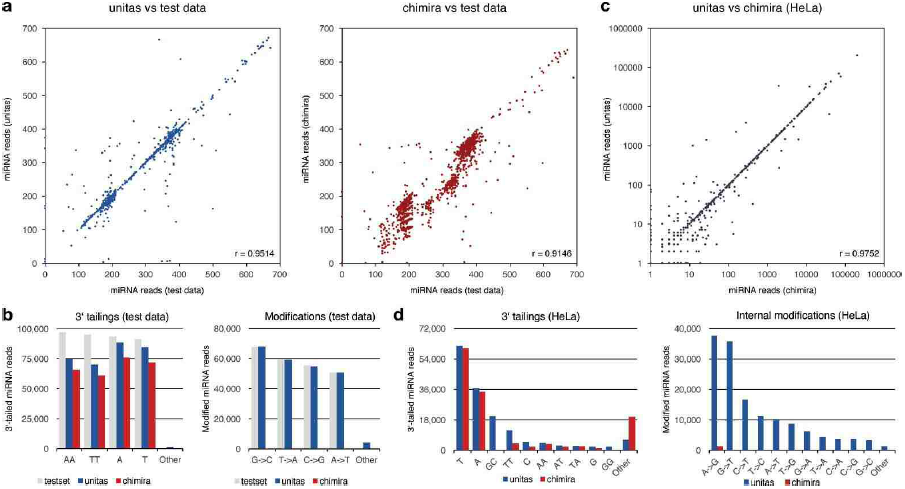
Analysis of miRNA expression and modification by unitas compared to Chimira. **a** Correlations of reads assigned to different miRNA genes by unitas and Chimira compared to reads profile of artificial dataset. For better comparison of miRNA counts, reads from same miRNA gene were combined. **b** 3’ tailings and internal modifications detected by unitas and Chimira in test data. **c** miRNA expression in HeLa cells determined by unitas compared to Chimira on logarithmic scale. **d** miRNA 3’ tailings and internal modifications quantified by unitas and Chimira in HeLa cells.

Subsequently, both tools were tested on published RNA-seq data obtained from HeLa cells (SRA accession: SRR029l24) [38]. The resulting miRNA expression profiles are highly similar, with a Pearson correlation coefficient of 0.9752 (Fig. 3c). While unitas found 96l,840 miRNA reads, Chimira called 960,880, which increased to 96l,563 if the option ‘-split counts from paralogs’ was selected. The amount of identified uridylation and adenylation events of 3’ ends were largely similar with a slight advantage on the side of unitas, but other tailing patterns were detected to a much lesser degree by Chimira (Fig. 3d). Analogous to the artificial test data, Chimira did not identify internal modifications to a comparable extent as unitas, apart from some amount of ADAR-dependent edits (A-to-G), which was also the most frequent modification detected by unitas.

Since both, unitas and similar computational approaches for miRNA annotation, rely on miRBase, it should be noted that the quality of database annotations in general vary among species, particularly those which are less well studied. Therefore, we point to existing tools designed specifically for the *de novo* prediction of miRNAs, such as CAP-miRSeq [35] and Oasis [36].

### Annotation of tRNA-derived small RNAs

Currently, there are three major tools specific to the identification of tRNA-derived sncRNAs [39]. The tRFfinder of the tRF2Cancer web server package [40] is restricted to the analysis of human samples and considers sequences with lengths between l4 and 32 nt only. This, however, poses a limitation for the detection of longer tRNA-derived sncRNAs like tRNA-halves. For instance, 56% of tRNA-derived reads are larger than 32 nt in a sncRNA dataset of seminal exosomes (SRA accession: SRRl2007l2) [41].

The second tool, tDRmapper [14], is a command line based set of Perl scripts, which identifies tRNA-derived RNAs (tDRs) from l4 to 40 nt in human and murine samples. Other species can be added manually according to a provided guide and with the help of Perl scripts that depend on bedtools. Notably, there are some features to the algorithm of tDRmapper that may hamper appropriate interpretation of the obtained results. First, sequences with l00 reads or less are discarded. Subsequently, so-called primary tDRs are determined by the location, at which more than 50% of all reads mapping to a source tRNA are aligned. For example, the 5’-tRF type is assigned if more than 50% map at the 5’-end of the source tRNA. Moreover, tRFs are defined by length as being smaller than 28 nt and tRNA-halves as 28 nt or larger, regardless of alignment position. Further, a primary tDR is only specified if more than 66% of all reads mapping to the source tRNA map to any position of the considered tDR. Lastly, tDRs are quantified by ‘relative abundance’, which is calculated by multiplying the percentage of tDR reads that map to its source tRNA and the proportion of reads on the area with the highest read coverage across the source-tRNA. lmportantly, the resulting counts are not normalized, meaning there is no fractional assignment for multi-mapping reads. As the authors themselves point out, this approach may overestimate the relative abundance of a primary tDR.

Finally, another command line based tool called MlNTmap was recently developed for the profiling of tRNA fragments from human small RNA-seq data, emphasizing the profiling of both nuclear and mitochondrial tRNA fragments [15]. However, tRFs generated from trailer sequences (tRF-l) and 5’ leader-tRFs are excluded from analysis.

To test the efficiency and accuracy of unitas in the detection of tRNA-derived small RNAs, we produced an artificial dataset based on human tRNAs from the genomic tRNA database, incorporating one mismatch in 50% of sequences. Running with default settings, unitas assigned 92% of reads to tRNAs, which increased to 97% if miRNA detection was skipped (option ‘- nomiR’). For running tDRmapper on the test data, we disabled the rejection of sequences with less than l0l reads, since this would eliminate the entire input. Both, tDRmapper and MlNTmap, deviated considerably from the original dataset in read shares assigned to different tRNA-derived sRNA classes, in contrast to unitas (Fig. 4a). A direct comparison of read counts, however, was not possible due to the non-normalizing quantification method of tDRmapper, which calculates so-called relative abundance. Further, read shares were most precisely assigned to source tRNAs by unitas, indicated by the highest Pearson correlation coefficient (0.9896) among the tested tools (Fig. 4b).

**Fig. 4:**
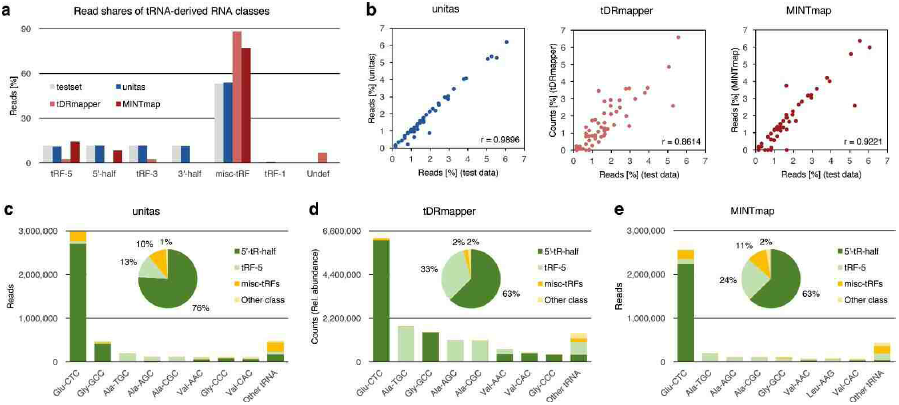
Detection of tRNA-derived small RNAs by unitas, tDRmapper and MlNTmap. **a** Read shares assigned to different tRNA-derived RNA classes of artificial test data by unitas and other tools compared to test set profile. Misc-tRFs (miscellaneous tRFs) are equivalent to internal tRFs. On test data, the elimination of sequences with less than l0l reads by tDRmapper was disabled. **b** Correlation of read proportions allocated to source tRNAs by unitas and other tools with original test data read shares per tRNA. **c** Analysis of RNA-seq data from seminal exosomes by unitas, **d** tDRmapper and **e** MlNTmap.

Additionally, unitas, tDRmapper and MlNTmap were tested on RNA-seq data of exosomes from seminal fluid (SRRl2007l2) [41], using default settings. The differences in results between unitas (Fig. 4c) and tDRmapper (Fig. 4d) are largely due to the lack of fractional assignment for multi-mapping reads in the quantification approach of tDRmapper. Analysis by MlNTmap yielded results that are largely similar to the output of unitas, but with overall slightly lower read counts and changed order of the source tRNAs with descending read coverage (Fig. 4e). Details on the test dataset and the annotation of tRFs from artificial and biological data with different tools are available in Additional file 5: Table S4 (A-G).

### Annotation of phasiRNAs

We tested and compared phasiRNA annotation performance of unitas and PhaseTank, which is currently the only published tool for prediction of phased RNAs [16]. We used artificial datasets with known amounts of phased RNAs (Additional file 6: Table S5) as well as biological small RNA data from panicles of the two rice strains 93-ll and Nipponbare [42] to predict phased RNAs with unitas and PhaseTank using default parameters. All test datasets were collapsed to non-identical sequences, retaining information on read counts for each sequence in the FASTA header.

Subsequently, the datasets were formatted to satisfy PhaseTank requirements (special format of FASTA headers) and used as input for PhaseTank (v.l.0) using default settings to search for 2l nt phased RNAs. Subsequently we searched for 24 nt phased RNAs with PhaseTank using the option ‘-size 24’. To generate input files for unitas, we mapped the test datasets to the human genome (GRCh38) with bowtiel, bowtie2 and STAR using settings that correspond to recommended and widely used settings for mapping of small RNAs with these tools and considering only perfect matches to be in line with PhaseTank default settings. The resulting SAM alignment files were used as input for unitas which was started twice with the option ‘-phasi 2l’ or ‘-phasi 24’, respectively, to search for 2l nt and 24 nt phased RNAs.

Using artificial datasets, we found that both tools perform equally well with those datasets that comprise exclusively phased RNAs or have rather low amounts of non-phased RNAs (Fig. 5a). However, PhaseTank drastically loses its sensitivity with an increasing amount of non-phased sequences within a dataset, while the sensitivity of unitas remains unaffected (Fig. 5a). When we assigned a read count value of l0 to each artificial phased RNA sequence, PhaseTank performs approximately as well as unitas, illustrating that sensitivity of PhaseTank not only depends on the number of phased sequences, but also on the number of reads per phased sequence. Consequently, PhaseTank will particularly misses those phasiRNA-producing loci that have a low sequence read coverage (Fig. 5a and 5b). For neither tool we observed a namable issue with false positive predicted phasiRNAs when running both programs with datasets comprising no phased RNAs (Additional file 7: Table S6).

**Fig. 5:**
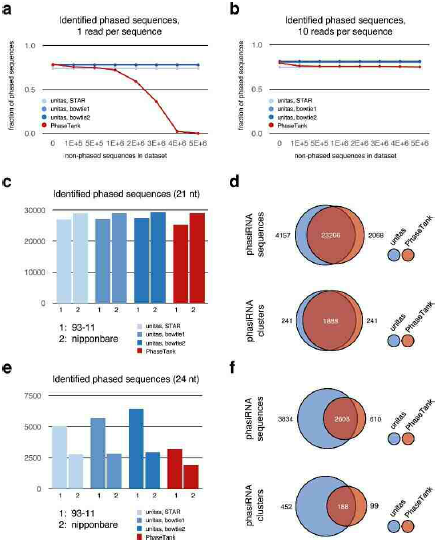
Identification of phased RNAs with unitas and PhaseTank. a ldentified phased RNAs in artificial datasets, with one read per phased RNA and growing amount of background sequences. b ldentified phased RNAs in artificial datasets, with ten reads per phased RNA and growing amount of background sequences. c Number of identified 2l nt phasiRNAs in small RNA datasets from rice panicles. d Congruency between unitas and PhaseTank results for 2l nt phasiRNA prediction. e Number of identified 24 nt phasiRNAs in small RNA datasets from rice panicles. f Congruency between unitas and PhaseTank results for 24 nt phasiRNA prediction.

When searching for 2l nt phased RNAs in biological datasets we noted that unitas identifies slightly more phasiRNAs compared to PhaseTank, while the number of identified clusters was identical (Fig. 5c and 5d). Overall, the congruency between unitas and PhaseTank results is very high (Fig. 5d). However, when searching for 24 nt phased RNAs in the same datasets we observed remarkable differences with unitas identifying both more phasiRNA sequences and more phasiRNA clusters (Fig. 5e and 5f). Considering that PhaseTank is less sensitive when the fraction of phased RNAs within a given dataset is low, these results are in line with the fact that the abundance of 24 nt phasiRNAs in rice panicles is several times lower compared to 2l nt phasiRNAs [42]. Accordingly, phasiRNA clusters identified only by unitas have relatively low read coverage, while phasiRNA clusters identified only by PhaseTank were rejected by unitas because of a high strand bias (> 95%) of mapped sequence reads.

### Complete annotation of NGS datasets

To emphasize the broad range of possible applications of unitas on diverse sncRNA-seq datasets, we analyzed three exemplary libraries, which differ in origin and structure (Fig. 6). The sncRNA annotation output of unitas provides a general overview of the small RNA composition, as shown for a dataset produced from HeLa cancer line cells (SRA accession: SRR029l24, Fig. 6a) [38]. ln this library, the largest fraction of sncRNAs is constituted by miRNAs, which are of growing interest for cancer studies and clinical trials using miRNA profiling for patient diagnosis [43]. Apart from expression profiles and 3’ tailings, unitas offers a convenient description of miRNA modifications per position (Fig. 6b). For target recognition, complementarity of the seed region of a miRNA (positions 2-7) to its target is critical for downstream silencing efficacy. Beyond the seed region, a strong sequence conservation can also be observed at position 8 [44], and finally, miRNA sequences frequently start with a uridine which was found to promote miRNA loading on Argonaute proteins [45]. According to these functional aspects, the unitas output shows that in HeLa cells the first eight positions from the 5’ end are rarely modified. lnternal modifications occur predominantly at distinct positions downstream of the seed region, with A-to-G (A- to-l), known as ADAR edits [46], and G-to-T being the most common, followed by modifications leading to uridine or guanine incorporation (Fig. 6b).

**Fig. 6:**
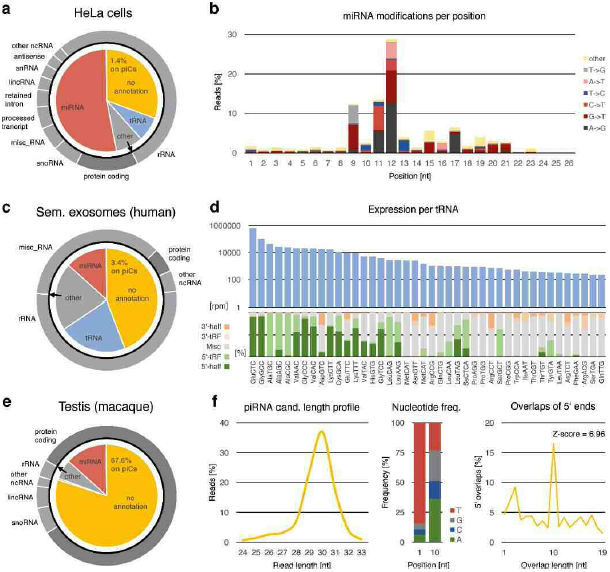
Exemplary analysis of three small RNA libraries by unitas. **a** General sncRNA annotation of sRNA- seq data from HeLa cells (SRR029l24). **b** Overall miRNA modification per position in HeLa cells data. **c** General sncRNA annotation of sRNA-seq data from exosomes of human seminal fluid (SRRl2007l2). **d** Read coverage of tRNAs as reads per million (rpm) on logarithmic scale and percentages of tRNA-derived sRNA classes per tRNA. **e** General sncRNA annotation of sRNA-seq data from macaque testis (SRR55358l). **f** Rates of 5’ overlaps and nucleotide frequencies of piRNA candidate reads, i.e. not annotated reads that map to known piRNA producing loci or piRNA clusters (piCs).

Next, we analyzed a library generated from exosomes of human seminal fluid (SRA accession: SRRl2007l2) [41]. Notably, this dataset is particularly abundant in tRNA-derived reads, while containing less miRNAs and other annotated reads (Fig. 6c). For the analysis of such sequences, unitas provides a summary of read counts for each tRNA gene, apportioned to classes of tRNA- derived sncRNAs. The majority of tRNA-derived reads in this library is represented by 5’ halves, originating mainly from the tRNAs Glu-CTC and Gly-GCC (Fig. 6d). lt has been shown that fragments specifically derived from 5’ ends of tRNA-Gly-GCC in mouse epididymosomes repress genes by regulating the endogenous retroelement MERVL, whereas an RNA interference against the middle or 3’ end of tRNA-Gly-GCC and other tRNAs had no effect on these MERVL-dependent genes [27]. Generally, we found 5’ halves and 5’ tRFs to be the dominant classes, being especially associated with tRNAs exhibiting high read coverage (Fig. 6d). With decreasing coverage, however, tRNAs tend to give rise primarily to internal fragments and to a lesser extent to 3’ halves and 3’ tRFs. This suggests that internal fragments, which we describe as miscellaneous tRFs (misc-tRFs), might be rather characterized as random debris of degraded tRNAs than a class of tRNA-derived small RNAs in its own right.

Lastly, we chose a sRNA dataset from macaque testis for analysis (SRA accession: SRR55358l) [47]. As expected, the vast majority of testis-expressed sRNAs did not match any class of known non-coding RNA (Fig. 6e). ln contrast to the former libraries, unitas found that the bulk of non-annotated reads (67.6%) maps to known piRNA producing loci. Besides the typical length profile (Fig. 6f), these piRNA candidate sequences show a strong bias for uridine at 5’ ends (84.5%) which is typical for primary piRNAs being processed by the endonuclease Zucchini (PLD-6) [48, 49]. Moreover, unitas attests a significant ping-pong signature (Z- score = 6.96), namely a high rate of l0 nt 5’ overlaps of sense and antisense reads, which is a hallmark of secondary piRNA biogenesis via the ping-pong cycle [50, 51, 52].

## Discussion

Small RNA biology has become a major field in molecular biology research. ln 20l6, sequence data from 727l lllumina sequencing runs with miRNA sequencing strategy, comprising more than 82 billion sequence reads, was uploaded to NCBl’s Sequence Read Archive. Assuming total costs of l5 USD per l Million clean sequence reads, the total miRNA sequencing value for 20l6 amounts to l.2 million USD. Noteworthy, these numbers only refer to published sequence data and certainly only mirror the tip of the iceberg. Nevertheless, in light of these numbers, even a seemingly trivial improvement of adapter recognition and trimming by unitas yields a surplus value of more than l00,000 USD per year, considering the amount of additionally mapped sequence reads. However, although this is a benefit of unitas that can be descriptively quantified, it clearly reflects only a minor aspect of the overall value of unitas.

Within the field of small RNA biology, miRNAs receive widespread attention owing to their pervasive contribution to gene regulatory processes [53]. However, it is not only mere miRNA expression, but also their post-transcriptional modification that vitally affects miRNA activity. Uridylation of miRNAs is thought to play a role in miRNA stability and possibly marks small RNAs for degradation [54, 55]. Adenylation has recently been linked to clearance of maternal miRNAs in *Drosophila* eggs [56]. Further, internal modification events of miRNAs (or their precursors) can have wide implications for miRNA biogenesis and function [57, 58, 59, 60]. lt is therefore of immense importance, to accurately identify post-transcriptional editing events to gain a deeper understanding of miRNA- dependent regulatory processes. As we have shown, unitas is more sensitive in detecting 3’ tailing events and much more sensitive in detecting internal modifications compared to existing tools. lmportantly, unitas not only focuses on well-known adenylation, uridylation and ADAR-dependent A-to-l editing, but also allows to detect all other types of modification events which can greatly facilitate the detection of yet unknown enzymatic editing activity in the future.

tRNA-derived small RNAs have been regarded as simple and non-functional degradation products for a long time. However, strong evidence for diverse functional roles in gene regulation, cancer biology, apoptosis and protein synthesis is mounting [24, 25, 26, 27, 28, 29, 30]. Since tRNAs and their precursor transcripts can be processed into functionally distinct types of tRFs, accurate attribution of tRNA-derived small RNAs to the different types of tRFs is important to make functional interpretations. ln this regard, unitas shows higher precision than existing tools, while being also more sensitive in overall tRF detection. Notably, the recently published tool for tRF annotation MlNTmap [15], which is more sensitive and accurate than the older tDRmapper [14] cannot identify some of the yet rather enigmatic tRFs which have their origin beyond the mature tRNA molecule, namely tRF-l and 5’ leader-tRFs. Here, unitas can enable researches to elucidate possible functions of these cryptic RNAs by first of all spotting them in sncRNA transcriptomes.

Initially described as trans-acting siRNAs [61], plant specific phasiRNAs are well-characterized actors in post-transcriptional gene silencing [62]. ln most cases, phasiRNAs are 2l nt in length, but different pathways that produce 22 nt and 24 nt phasiRNAs have been described as well. Since the latter are by far less abundant, their detection in small RNA transcriptomes is challenging and the current approach underestimates the number of, e.g., phased 24 nt RNAs in small RNA datasets from rice panicles. ln contrast, sensitivity of unitas depends far less on the amount of background reads, making it more suitable for the detection of particularly low abundance phasiRNAs and their source loci.

## Conclusions

So far, accurate annotation of sncRNA required a large set of different software tools with a number of additional prerequisites. While the installation of these bioinformatics tools can pose a challenge for the non-expert user, unitas brings together all annotation and analysis features with an absolute minimum of further requirements. A complete annotation run is finished within a few minutes and can be started with one simple shell command, making its usage very convenient. By facilitating sncRNA annotation and providing in depth analyses that previously was not accessible for the non-expert user, we believe that unitas is a valuable tool for researchers in all fields connected to small RNA biology.

## Declarations

### Acknowledgements

We thank Hans Zischler, Hendrik Dorschmann, Sabrina Saurin, Julia Jehn and Julian Kiefer for testing unitas in the course of their projects and providing valuable feedback how to improve this software

### Authors’ contributions

DR has designed the unitas software, DG and DR coded it. The software was tested by CH, DG and DR. ln-depth analysis of existing annotation algorithms was done by CH and DG, Figures were designed by DG and DR. The manuscript was written by DG and DR with valuable input of CH. All authors read and approved the final manuscript.

### Availability of data and material

The unitas software and a detailed documentation are freely available at https://sourceforge.net/projects/unitas/ and at http://www.smallrnagroup.uni-mainz.de/software.html. Additional Perl scripts, test datasets and exemplary unitas output files are available at http://www.smallrnagroup.uni-mainz.de/data/UNlTAS/resources.html.

### Ethics approval and consent to participate

This study required no ethics approval.

### Consent for publication

This article is distributed under the terms of the Creative Commons Attribution 4.0 lnternational License (http://creativecommons.org/licenses/by/4.0/), which permits unrestricted use, distribution, and reproduction in any medium, provided you give appropriate credit to the original authors and the source, provide a link to the Creative Commons license, and indicate if changes were made. The Creative Commons Public Domain Dedication waiver (http://creativecommons.org/publicdomain/zero/l.0/) applies to the data made available in this article, unless otherwise stated

### Additional Material

**Additional file 1.** File format: .pdf. Title of data: Supplememtary Methods. Description of data: Describes how the artificial sncRNA datasets that were used for testing purposes were generated.

**Additional file 2.** File format: .xlsx. Title of data: Table Sl. Description of data: Detailed results for adapter prediction and trimming.

**Additional file 3.** File format: .xlsx. Title of data: Table S2. Description of data: Detailed results of low complexity filtering.

**Additional file 4.** File format: .xlsx. Title of data: Table S3. Description of data: Detailed results of miRNA annotation.

**Additional file 5.** File format: .xlsx. Title of data: Table S4. Description of data: Detailed results of tRNA annotation.

**Additional file 6.** File format: .xlsx. Title of data: Table S5. Description of data: lnformation on artificial and semi-artificial test datasets for phasiRNA prediction.

**Additional file 7.** File format: .xlsx. Title of data: Table S6. Description of data: Detailed results of phasiRNA prediction.

**Additional file 8.** File format: .xlsx. Title of data: Table S7. Description of data: Detailed description of the artificial miRNA test dataset.

## References

1. Holoch D, Moazed D. RNA-mediated epigenetic regulation of gene expression. Nat Rev Genet. 20l5;l6:7l–84.

2. Gebert D, Rosenkranz D. RNA-based regulation of transposon expression. WlRES RNA 20l5;6:687–708.

3. Gorski SA, Vogel J, Doudna JA. RNA-based recognition and targeting: sowing the seeds of specificity. Nat Rev Mol Cell Biol. 20l7;l8:2l5–228.

4. Li Z, Ender C, Meister G, Moore PS, Chang Y, John B. Extensive terminal and asymmetric processing of small RNAs from rRNAs, snoRNAs, snRNAs, and tRNAs. Nucleic Acids Res. 20l2;40:6787–6799.

5. Daugaard l, Hansen TB. Biogenesis and Function of Ago- Associated RNAs. Trends Genet. 20l7;33:208–2l9.

6. Sarkies P, Miska EA. Small RNAs break out: the molecular cell biology of mobile small RNAs. Nat Rev Mol Cell Biol. 20l4;l5:525–535.

7. Rashed MH, Bayraktar E, Helal GK, Abd-Ellah MF, Amero P, Chavez-Reyes A, et al. Exosomes: From Garbage Bins to Promising Therapeutic Targets. lnt J Mol Sci. 20l7;l8:E538.

8. Davis MPA, van Dongen S, Abreu-Goodger C, Bartonicek N, Enright AJ. Kraken: A set of tools for quality control and analysis of high- throughput sequence data. Methods. 20l3;63:4l–49.

9. Tsuji J, Weng Z. DNApi: A de novo adapter prediction algorithm for small RNA sequencing data. PLoS One. 20l6;ll:e0l64228.

10. Martin M. Cutadapt removes adapter sequences from high-throughput sequencing reads. EMBnet.journal. 20ll;l7:0–12.

11. Morgulis A, Gertz EM, Schaffer AA, Agarwala R. A fast and symmetric DUST implementation to mask low-complexity DNA sequences. J Comput Biol. 2006;l3:028–1040.

12. Dozmorov MG, Adrianto l, Giles CB, Glass E, Glenn SB, Montgomery C, et al. Detrimental effects of duplicate reads and low complexity regions on RNA- and ChlP-seq data. BMC Bioinformatics. 20l5;l6(Suppl l3):Sl0.

13. Vitsios DM, Enright AJ. Chimira: Analysis of small RNA sequencing data and microRNA modifications. Bioinformatics. 20l5;3l:3365–3367.

14. Selitsky SR, Sethupathy P. tDRmapper: challenges and solutions to mapping, naming, and quantifying tRNA-derived RNAs from human small RNA-sequencing data. BMC Bioinformatics. 20l5;l6:354.

15. Loher P, Telonis AG, Rigoutsos l. MlNTmap: fast and exhaustive profiling of nuclear and mitochondrial tRNA fragments from short RNA-seq data. Sci Rep. 20l7;7:4ll84.

16. Guo Q, Qu X, Jin W. PhaseTank: genome-wide computational identification of phasiRNAs and their regulatory cascades. Bioinformatics. 20l5;3l:284–286.

17. Rosenkranz D. piRNA cluster database: a web resource for piRNA producing loci. Nucleic Acids Res. 20l6;44:D223–230.

18. Aken BL, Achuthan P, Akanni W, Amode MR, Bernsdorff F, Bhai J, et al. Ensembl 20l7. Nucleic Acids Res. 20l7;45:D635–D642.

19. Kozomara A, Griffiths-Jones S. miRBase: annotating high confidence microRNAs using deep sequencing data. Nucleic Acids Res. 20l4;42:D68–73.

20. Chan PP, Lowe TM. GtRNAdb 2.0: an expanded database of transfer RNA genes identified in complete and draft genomes. Nucleic Acids Res. 20l6;44:Dl84–l89.

21. Quast C, Pruesse E, Yilmaz P, Gerken J, Schweer T, Yarza P, et al. The SlLVA ribosomal RNA gene database project: improved data processing and web-based tools. Nucleic Acids Res. 20l3;4l:D590–596.

22. Rosenkranz D, Han CT, Roovers EF, Zischler H, Ketting RF. Piwi proteins and piRNAs in mammalian oocytes and early embryos: From sample to sequence. Genom Data. 20l5;5:309–3l3.

23. Jiang H, Wong WH. SeqMap: mapping massive amount of oligonucleotides to the genome. Bioinformatics. 2008;24:2395–2396.

24. Lee YS, Shibata Y, Malhotra A, Dutta A. A novel class of small RNAs: tRNA-derived RNA fragments (tRFs). Genes Dev. 2009;23:2639–2649.

25. Sobala A, Hutvagner G. Small RNAs derived from the 5’ end of tRNA can inhibit protein translation in human cells. RNA Biol. 20l3;l0:553–563.

26. Keam SP, Hutvagner G. tRNA-Derived Fragments (tRFs): Emerging New Roles for an Ancient RNA in the Regulation of Gene Expression. Life. 20l5;5:638–1651.

27. Sharma U, Conine CC, Shea JM, Boskovic A, Derr AG, Bing XY, et al. Biogenesis and function of tRNA fragments during sperm maturation and fertilization in mammals. Science. 20l6;35l:396–39l.

28. Venkatesh T, Suresh PS, Tsutsumi R. tRFs: miRNAs in disguise. Gene. 20l6579:33–l38.

29. Kumar P, Kuscu Cl, Dutta A. Biogenesis and Function of Transfer RNA-Related Fragments (tRFs). Trends Biochem Sci. 20l6;4l:679–689.

30. Keam SP, Sobala A, Ten Have S, Hutvagner G. tRNA-Derived RNA Fragments Associate with Human Multisynthetase Complex (MSC) and Modulate Ribosomal Protein Translation. J Proteome Res. 20l7;l6:4l3–420.

31. Zhang Z, Xu J, Koppetsch BS, Wang J, Tipping C, Ma S, et al. Heterotypic piRNA Ping-Pong requires qin, a protein with both E3 ligase and Tudor domains. Mol Cell. 20ll;44:572–584.

32. Howell MD, Fahlgren N, Chapman EJ, Cumbie JS, Sullivan CM, Givan SA, et al. Genome-wide analysis of the RNA-DEPENDENT RNA POLYMERASE6/DlCER-LlKE4 pathway inArabidopsis reveals dependency on miRNA-and tasiRNA-directed targeting. Plant Cell. 2007;l9:926–942.

33. Schmitt AO, Herzel H. Estimating the entropy of DNA sequences. J Theor Biol. 1997;l88:369–377.

34. Wootton JC, Federhen S. Analysis of compositionally biased regions in sequence databases. Methods Enzymol. 1996;266:554–57l.

35. Sun Z, Evans J, Bhagwate A, Middha S, Bockol M, Yan H, et al. CAP-miRSeq: a comprehensive analysis pipeline for microRNA sequencing data. BMC Genomics. 20l4;l5:423.

36. Capece V, Garcia Vizcaino JC, Vidal R, Rahman RU, Pena Centeno T, Shomroni O, et al. Oasis: online analysis of small RNA deep sequencing data. Bioinformatics. 20l5;3l:2205–2207.

37. Zheng Y, Ji B, Song R, Wang S, Li T, Zhang X, et al. Accurate detection for a wide range of mutation and editing sites of microRNAs from small RNA high-throughput sequencing profiles. Nucleic Acids Res. 20l6;44:el23.

38. Mayr C, Bartel DP. Widespread Shortening of 3’UTRs by Alternative Cleavage and Polyadenylation Activates Oncogenes in Cancer Cells. Cell. 2009;l38:673–684.

39. Xu W-L, Yang Y, Wang Y-D, Qu L-H, Zheng L-L. Computational Approaches to tRNA-Derived Small RNAs. Non-Coding RNA. 20l7;3:2.

40. Zheng L, Xu W, Liu S, Sun W, Li J, Wu J, et al. tRF2Cancer: A web server to detect tRNA-derived small RNA fragments (tRFs) and their expression in multiple cancers. Nucleic Acids Res. 20l6;44:Wl85–l93.

41. Vojtech L, Woo S, Hughes S, Levy C, Ballweber L, Sauteraud RP, et al. Exosomes in human semen carry a distinctive repertoire of small non-coding RNAs with potential regulatory functions. Nucleic Acids Res. 20l4;42:7290–7304.

42. Song X, Li P, Zhai J, Zhou M, Ma L, Liu B, et al. Roles of DCL4 and DCL3b in rice phased small RNA biogenesis. Plant J. 20l2;69:462–474.

43. Hayes J, Peruzzi PP, Lawler S. MicroRNAs in cancer: Biomarkers, functions and therapy. Trends Mol Med. 20l4;20:460–469.

44. Lewis BP, Burge CB, Bartel DP. Conserved seed pairing, often flanked by adenosines, indicates that thousands of human genes are microRNA targets. Cell. 2005;1205–20.

45. Seitz H, Tushir JS, Zamore PD. A 5’-uridine amplifies miRNA/miRNA* asymmetry inDrosophila by promoting RNA-induced silencing complex formation. Silence. 20ll;2:4.

46. Kawahara Y, Zinshteyn B, Sethupathy P, lizasa H, Hatzigeorgiou AG, Nishikura K. Redirection of silencing targets by adenosine-to-inosine editing of miRNAs. Science. 2007;3l5:l37–1140.

47. Meunier J, Lemoine F, Soumillon M, Liechti A, Weier M, Guschanski K, et al. Birth and expression evolution of mammalian microRNA genes. Genome Res. 20l3;23:34–45.

48. lpsaro JJ, Haase AD, Knott SR, Joshua-Tor L, Hannon GJ. The structural biochemistry of Zucchini implicates it as a nuclease in piRNA biogenesis. Nature. 20l2;49l:279–283.

49. Nishimasu H, lshizu H, Saito K, Fukuhara S, Kamatani MK, Bonnefond L, et al. Structure and function of Zucchini endoribonuclease in piRNA biogenesis. Nature. 20l2;49l:284–287.

50. Brennecke J, Aravin AA, Stark A, Dus M, Kellis M, Sachidanandam R, et al. Discrete small RNA-generating loci as master regulators of transposon activity inDrosophila. Cell. 2007;l28:089–1103.

51. Gunawardane LS, Saito K, Nishida KM, Miyoshi K, Kawamura Y, Nagami T, et al. A slicer-mediated mechanism for repeat-associated siRNA 5’ end formation in Drosophila. Science. 2007;3l5:587–l590.

52. Aravin AA, Sachidanandam R, Bourc’his D, Schaefer C, Pezic D, Toth KF, et al. A piRNA pathway primed by individual transposons is linked to de novo DNA methylation in mice. Mol Cell. 2008;3l:785–799.

53. Jonas S, lzaurralde E. Towards a molecular understanding of microRNA-mediated gene silencing. Nat Rev Genet. 20l5;l6:42l–433.

54. Li J, Yang Z, Yu B, Liu J, Chen X. Methylation protects miRNAs and siRNAs from a 3’-end uridylation activity inArabidopsis. Curr Biol. 2005;l5:50l–1507.

55. Ameres SL, Horwich MD, Hung JH, Xu J, Ghildiyal M, Weng Z, et al. Target RNA-directed trimming and tailing of small silencing RNAs. Science. 20l0;328:534–1539.

56. Lee M, Choi Y, Kim K, Jin H, Lim J, Nguyen TA, et al. Adenylation of maternally inherited microRNAs by Wispy. Mol Cell. 20l4;56:696–707.

57. Nishikura K. Functions and regulation of RNA editing by ADAR deaminases. Annu Rev Biochem. 20l0;79:321–349.

58. Chawla G, Sokol NS. ADAR mediates differential expression of polycistronic microRNAs. Nucleic Acids Res. 20l4;42:5245–5255.

59. Cui Y, Huang T, Zhang X. RNA editing of microRNA prevents RNA-induced silencing complex recognition of target mRNA. Open Biol. 20l5;5:50l26.

60. Behm M, Ohman M. RNA Editing: A Contributor to Neuronal Dynamics in the Mammalian Brain. Trends Genet. 20l6;32:65–l75.

61. Vazquez F, Vaucheret H, Rajagopalan R, Lepers C, Gasciolli V, Mallory AC, et al. Endogenous trans-acting siRNAs regulate the accumulation of Arabidopsis mRNAs. Mol Cell. 2004;l6:69–79.

62. Fei Q, Xia R, Meyers BC. Phased, secondary, small interfering RNAs in posttranscriptional regulatory networks. Plant Cell. 20l3;25:2400–24l5.

